# Reconstitution of autophagosomal membrane tethering reveals that Atg11 can bind and cluster vesicles on cargo mimetics

**DOI:** 10.1101/2023.12.19.572332

**Authors:** Devika Andhare, Sarah Katzenell, Sarah I Najera, Katherine M Bauer, Michael J Ragusa

## Abstract

Autophagy is essential for the degradation of mitochondria from yeast to humans. Mitochondrial autophagy in yeast is initiated when the selective autophagy scaffolding protein Atg11 is recruited to mitochondria through its interaction with the selective autophagy receptor Atg32. This also results in the recruitment of small 30 nm vesicles that fuse to generate the initial autophagosomal membrane. We demonstrate that Atg11 can bind to autophagosomal-like membranes in vitro in a curvature dependent manner via a predicted amphipathic helix. Deletion of the amphipathic helix from Atg11 results in a delay in the formation of mitophagy initiation sites in yeast. Furthermore, using a novel biochemical approach we demonstrate that the interaction between Atg11 and Atg32 results in the tethering of autophagosomal-like vesicles in clusters to giant unilamellar vesicles containing a lipid composition designed to mimic the outer mitochondrial membrane. Taken together our results demonstrate an important role for autophagosomal membrane binding by Atg11 in the initiation of mitochondrial autophagy.

## INTRODUCTION

Macroautophagy, hereafter autophagy, is the cellular process in which double-membrane vesicles, termed autophagosomes, are generated around cytosolic material leading to their capture within these vesicles^1^. Completed autophagosomes fuse with the vacuole, in yeast, or lysosomes, in higher eukaryotes, leading to the degradation of the captured contents. Autophagy can capture and degrade a wide range of cytosolic material including protein aggregates, intracellular pathogens and membrane-bound organelles. As such, autophagy is essential to maintain cellular homeostasis and also serves as an integral component of innate immunity^2,3^.

Cargo selection during autophagy can occur via a non-selective or selective mechanism^4^. In selective autophagy, cargos are marked as targets for autophagy by selective autophagy receptors (SARs)^5^. SARs are either soluble cytosolic proteins that commonly recognize ubiquitin moieties on the cargo to be degraded or, in the case of membrane-bound organelles, they can also be integral membrane proteins on the organelle to be degraded. To initiate selective autophagy, SARs recruit the selective autophagy scaffolding protein Atg11, in yeast, or FIP200, in higher eukaryotes, to the cargo to be degraded^6^. In yeast, Atg11 also binds to Atg9, an integral membrane protein embedded in 30 nm vesicles that serve as the initial membrane source for autophagy^7–9^. Once recruited to the cargo, Atg9 vesicles fuse to generate an initial membrane sheet, termed the phagophore^10^. The phagophore expands around the cargo through lipid transport from Atg2 in addition to other mechanisms^11–13^. Once the autophagosomal membrane has fully surrounded the cargo, it is closed leading to the capture of the marked cargo in a double membrane autophagosome^14^.

One of first critical steps in selective autophagy is the recruitment of Atg9 vesicles by Atg11 to autophagic cargo. Deletion of Atg11 in yeast was initially shown to block the recruitment of Atg9 to autophagy cargos highlighting the important role of Atg11 in recruiting membrane to cargo during selective autophagy initiation^15^. Atg11 was later shown to be required for the recruitment of Atg9 vesicles to cargo mimetic beads in vitro using biochemical reconstitution^9^. Following on this work, Atg11 was shown to directly interact with the flexible N-terminus of Atg9 via two PLF motifs^8^. More recently, truncations or mutations within the N-terminal domain of Atg11 have been shown to lead to a loss in the interaction between Atg11 and Atg9 in yeast^16^. These studies all point towards an essential role for Atg11 in the recruitment of membrane to cargos, specifically through the interaction between Atg11 and Atg9. However, while mutation of both PLF motifs in Atg9 led to a near complete loss in the interaction between Atg11 and Atg9, it did not fully prevent the recruitment of Atg9 to selective autophagy cargos^8^. Furthermore, the mammalian analog of Atg11, FIP200, can bind membranes directly, although the exact function of membrane binding by FIP200 is unknown^17^. Given these observations, we wanted to test if Atg11 is capable of binding membranes and whether this plays a role in selective autophagy.

## RESULTS

### Atg11 binds autophagosomal membranes in a clustered manner

To test if Atg11 can bind membranes we first purified Atg11 and GFP-Atg11 using Freestyle 293 cells as a recombinant expression system (**Figure S1**). Size exclusion chromatography of Atg11 in buffer containing 300mM NaCl had a similar elution profile and estimated molecular weight to previously published purifications of Atg11 from SF9 insect cells where it was shown to be a dimer in solution (**Figure S1B**)^18^. Next, we tested the ability of Atg11 to bind membranes. Both autophagosomal membranes and Atg9 vesicles are known to be composed of an unusually high concentration of negatively charged lipids, including between 37% and 44% of the negatively charged phosphatidylinositol (PI)^9,19^. Previously we have shown that several autophagy proteins including Atg23 and Atg16 bind membranes generated from yeast polar lipids (YPL), which also contain a high concentration of negatively charged lipids^20,21^. Therefore, we first tested if Atg11 can bind giant unilamellar vesicles (GUVs) made from YPL or a lipid mixture designed to mimic the autophagosomal membrane composition (43.9% phosphatidylcholine (PC), 19% phosphatidylethanolamine (PE), 32% phosphatidylinositol (PI), 5% phosphatidylinositol-3- phosphate (PI3P), and 0.1% rhodamine PE (RhPE) (automix+PI3P))^19^. 5% PI3P was added at the expense of PI as autophagosomal membranes contain PI3P but the exact concentration is unknown. GFP-Atg11 showed robust binding to YPL GUVs but was unable to bind autophagosomal membrane GUVs (**Figure S2A and B**). Given that Atg9 vesicles are highly curved while GUVs are essentially flat relative to the size of Atg11, we next investigated if curvature impacts the ability of Atg11 to bind membranes. For this we employed supported membrane templates (SMrT) that consist of an array of highly curved membrane tubes ranging from 20-60 nm in diameter and supported lipid bilayers (SLB) resting on a PEGylated glass coverslip^22,23^. The size of the SMrT nanotubes match the size of Atg9 vesicles that are reported to be 30-60 nm in diameter while SLBs are primarily flat membranes^10,24^. We made SMrT tubes and SLBs containing 43.9% PC, 19% PE and either 37% PI alone (automix) or 32% PI with 5% PI3P (automix+PI3P). Interestingly, GFP- Atg11 bound autophagosomal-like membranes as discreet clusters (**Figure 1A**). The presence of PI3P in membranes enhanced GFP-Atg11 recruitment. Importantly, the quantification of the ratio of GFP to membrane fluorescence showed that Atg11 binding was 5-fold higher on tubes (mean = 0.90 ± 0.4) than on SLB (mean = 0.16 ± 0.07) suggesting that Atg11 membrane binding is curvature sensitive (**Figure 1B**). As an independent validation of membrane binding, we also performed a floatation assay with automix and automix+PI3P liposomes. Atg11 migrated to the top layer of the sucrose density gradient in the presence of 100 nm diameter but not 1 µm diameter automix+PI3P liposomes further supporting the curvature sensitivity of Atg11 membrane binding (**Figure S3A**). DLS analysis also demonstrated that Atg11 binding to 100 nm liposomes increased the diameter of these vesicles from 107 nm to 260 nm suggesting that Atg11 may also be able to tether automix+PI3P liposomes (**Figure S3B**). These results suggest that Atg11 can bind autophagosomal-like membranes in a curvature dependent manner and that this binding is enhanced by the presence of PI3P.

**Figure 1.**
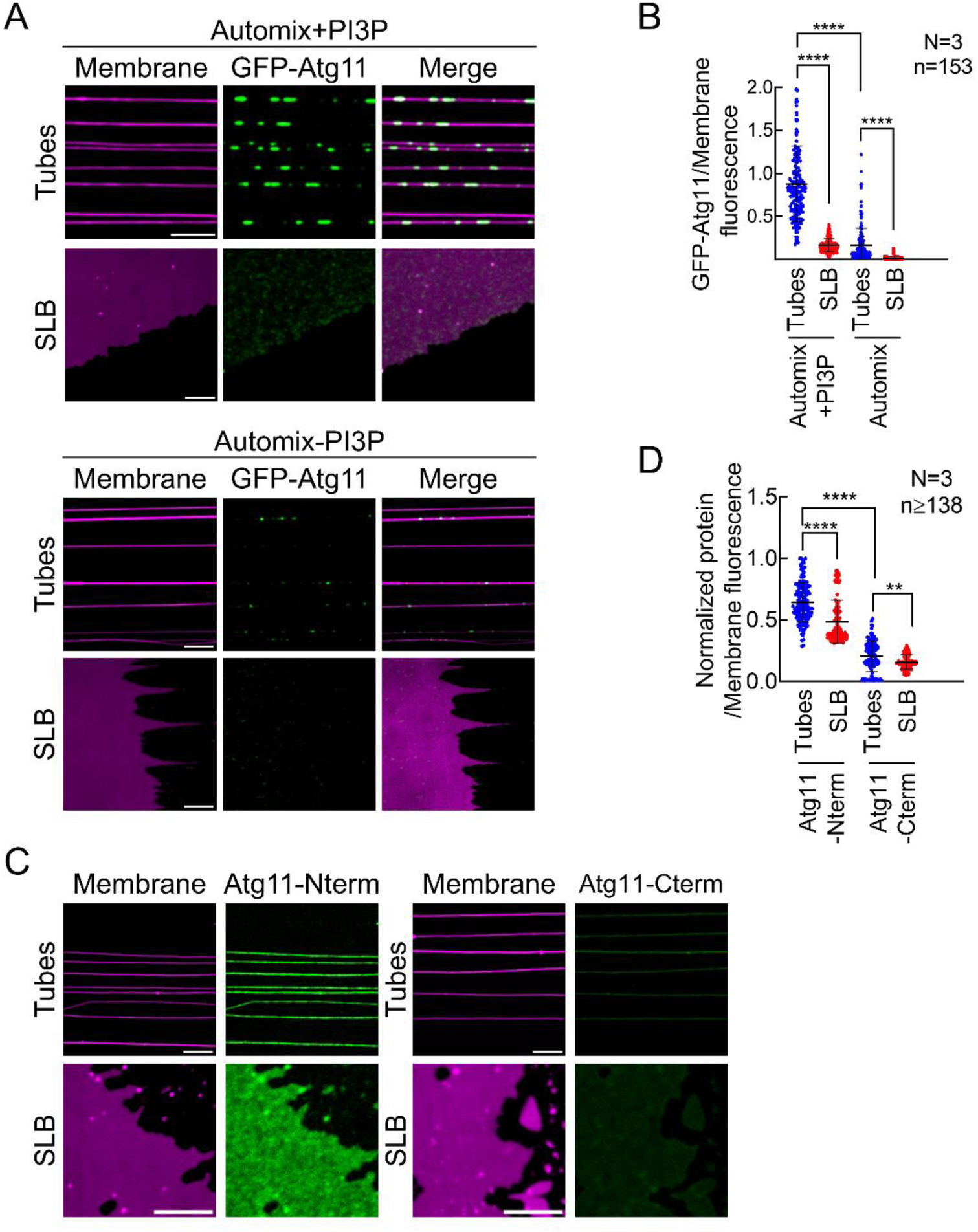
Atg11 binds curved autophagosomal-like membranes. (**A**) Representative images showing binding of GFP-Atg11 on an array of SMrT tubes and flat supported lipid bilayers (SLB). Membranes (magenta) are made of autophagosomal-like lipid composition with PI3P (Automix+PI3P, top panel) and without PI3P (Automix, bottom panel). Scale bar=5μm. (**B**) Quantification of GFP-Atg11 fluorescence normalized to the membrane fluorescence from SMrT tubes and SLB made with automix+PI3P and automix. A total 153 tubes or 2 to 4um^2^ individual patches of SLB (n=153) were analyzed across three independent repeats (N=3). Data represent mean ± SD. **** indicates p<0.0001 calculated using one-way ANOVA with Šídák’s multiple comparisons test. (**C**) Representative fluorescence images showing the binding of GFP-Atg11-Nterm and GFP-Atg11-Cterm on an array of tubes and SLB made of automix+PI3P. Scale bar=5μm. (**D**) Quantification of GFP-Atg11-Nterm or -Cterm normalized to membrane fluorescence for SMrT tubes and SLB made with automix+PI3P. A total of 138 to 150 tubes or 2 to 4um^2^ individual patches of SLB (n≥138) were analyzed across three independent repeats (N=3). Data represent mean ± SD. **** indicates p<0.0001 and ** indicates p=0.0086 calculated using one-way ANOVA with Šídák’s multiple comparisons test. Autophagosomal-like membranes with PI3P (automix+PI3P) were made of 43.9mol% PC, 19mol% PE, 32mol% PI, 5mol% PI3P with 0.1mol% RhPE. Autophagosomal-like membranes without PI3P (automix) were made similarly except for the omission of PI3P and increase of PI to 37mol%.

The N-terminal (amino acids 1-646) and C-terminal (amino acids 700-1178) regions of Atg11 can be purified using *E. coli* as a recombinant expression system^18^. Structure prediction of the N-terminal region of Atg11 (Atg11-Nterm) using AlphaFold 3 demonstrates that it forms a helical dimer with a structure that is similar to the N-terminal domain of FIP200 except that it is predicted to form a Z shaped dimer instead of a C shaped dimer (**Figure S4A**)^25,26^. The C-terminal region of Atg11 (Atg11-Cterm) contains a long coiled-coil domain, connected by linker regions to the Claw domain that binds SARs^27,28^. We purified GFP-tagged Atg11-Nterm and -Cterm and analyzed their membrane binding abilities on SMrT tubes and SLBs (**Figure 1C**). Quantitative fluorescence microscopy showed that Atg11-Nterm had significantly higher membrane binding than Atg11-Cterm suggesting that Atg11-Nterm contains the primary membrane binding region (**Figure 1D**). Moreover, unlike the full-length protein, Atg11-Nterm did not cluster on membranes and instead bound both membrane tubes and SLBs in a diffuse manner suggesting that membrane binding and clustering requires distinct regions of Atg11.

### Atg11 contains an amphipathic helix required for membrane binding

Sequence and structural analysis of the Atg11-Nterm revealed that it contains a predicted amphipathic helix spanning residues 612-646, with the highest hydrophobic moment (µH=0.546) occurring between residues 626 and 643 (**Figure 2A**). Amphipathic helices are known to mediate curvature- and charge-dependent membrane binding^29,30^. Importantly, this region in the predicted Atg11-Nterm structure has low and very low pLDDT scores, indicating that it is likely not in the correct conformation in the structure prediction (**Figure S4B**). This suggests that this helix may instead be available for membrane binding. Deletion of amino acids 612-646 from the Atg11-Nterm sequence and rerunning the AlphaFold3 prediction did not alter the overall dimer structure further suggesting that the predicted amphipathic helix is not required for formation of the Atg11-Nterm dimer structure (**Figure S4C**). We therefore deleted the predicted amphipathic helix in Atg11-Nterm (Atg11-Nterm^Δ612–646^) and purified it using *E. coli* as a recombinant expression system. We compared the elution profile of Atg11-Nterm^Δ612–646^ from size exclusion chromatography with that of Atg11-Nterm to verify that deletion of this helix did not disrupt the overall shape of the protein (**Figure S4D**). The ability of GFP-Atg11-Nterm^Δ612–646^ to bind membrane tubes was tested using SMrT tubes (**Figure 2B**). Comparison of GFP to membrane fluorescence ratios revealed that Atg11-Nterm^Δ612-646^ had 2.6-fold lower binding to membrane tubes than Atg11-Nterm (**Figure 2C**). Together these results suggest that Atg11 binds autophagosomal-like membranes primarily via an amphipathic helix between amino acids 612-646.

**Figure 2.**
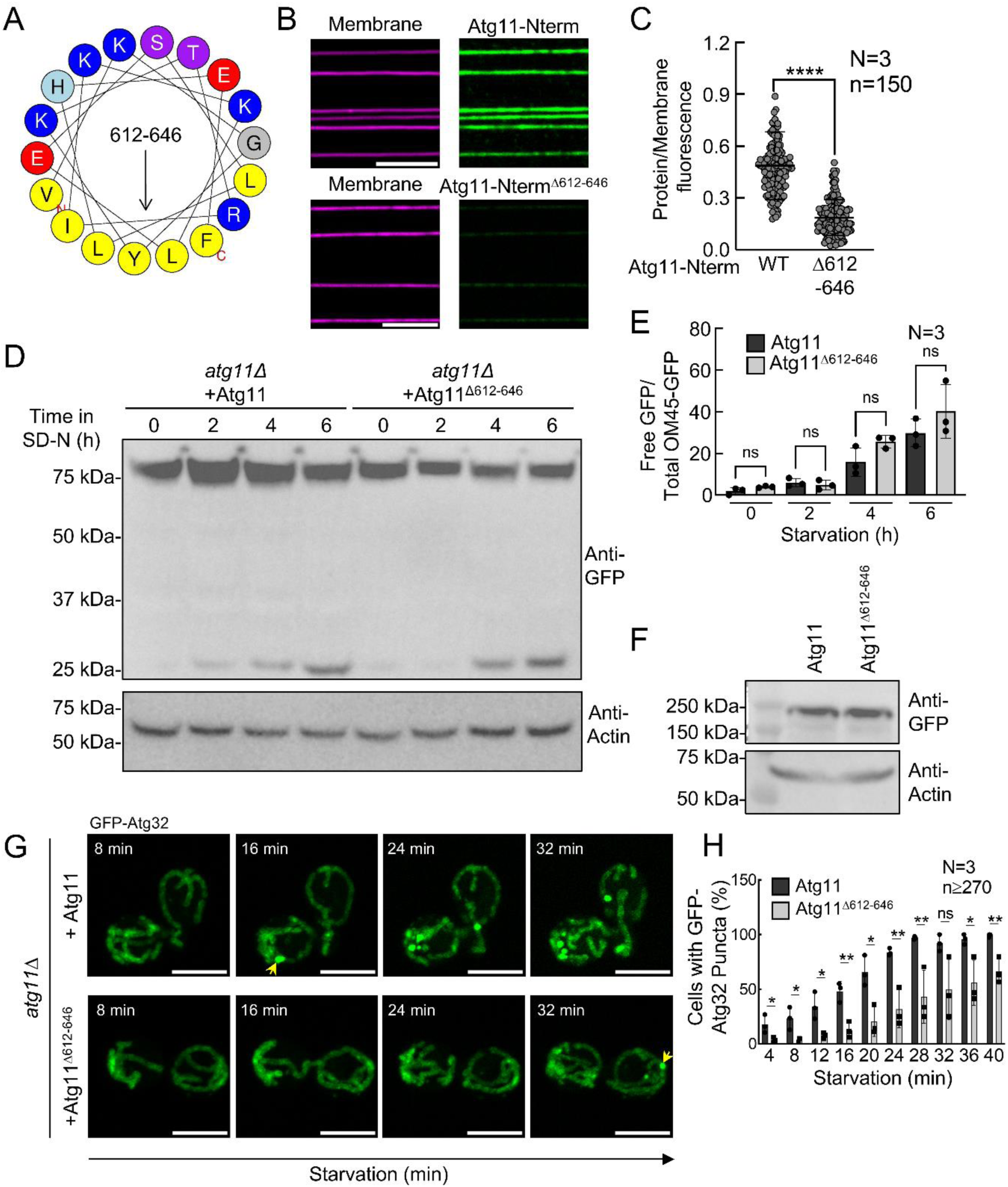
A putative amphipathic helix mediates autophagosomal membrane binding by Atg11 and contributes to mitophagy. (**A**) Helical wheel representation of the predicted amphipathic helix in Atg11-Nterm. (**B**) Representative images showing GFP-Atg11-Nterm or GFP-Atg11-Nterm^Δ612–646^ binding on SMrT tubes made with 43.9% PC, 19% PE, 32% PI, 5% PI3P with 0.1mol% RhPE (automix+PI3P). Scale bar=5μm. **C**) Quantification of GFP-Atg11-Nterm or GFP-Atg11-Nterm^Δ612–646^ fluorescence normalized by membrane fluorescence from **B**. Data represent mean ± SD of 150 tubes (n=150) analyzed across three independent repeats (N=3). **** indicates p<0.0001 calculated using the non-parametric Mann-Whitney’s test. (**D**) *atg11*Δ cells transformed with Atg11 or Atg11^Δ612–646^ were grown in media containing lactate and shifted to nitrogen starvation media (SD-N) to induce mitophagy. Western blot showing OM45-GFP at 0, 2, 4 and 6 hours after nitrogen starvation. Actin was used as a loading control. (**E**) Quantification of the background subtracted free GFP band normalized to total OM45-GFP at 0, 2, 4 and 6 hours after nitrogen starvation. Data represent mean ± SD from three independent repeats (N=3). ns indicates non-significant analyzed using one-way ANOVA with Šídák’s multiple comparisons test. **(F)** Western blot showing the expression of 2xGFP-Atg11 and 2xGFP-Atg11^Δ612–646^. (**G**) Timelapse fluorescence images showing *atg11Δ* cells expressing GFP-Atg32 with Atg11 (top panel) or Atg11^Δ612–646^ (bottom panel). Cells grown in nutrient rich (SMD) media were attached to a concanavalin A-treated coverslip, assembled in a flow cell maintained at 30°C. Nitrogen-starvation medium (SD-N) was introduced under controlled laminar flow, and time-lapse images were acquired three minutes after the flow initiation. Images were captured at 4 minutes interval. Yellow arrowheads indicate the onset of GFP-Atg32 puncta formation. Scale bar=5μm. (**H**) Quantification of cells subjected to SD-N showing percent of cells with at least one GFP-Atg32 punctum. Data represents mean ± SD of a total of 270 cells for Atg11 and 290 cells for Atg11^Δ612– 646^ (n≥270) from three independent repeats (N=3). * indicates p=0.0151 to p=0.0487, ** indicates p=0.0089, ns indicates non-significant calculated using multiple t tests.

To determine if the amphipathic helix of Atg11 is required for selective autophagy in cells we monitored the selective autophagy of mitochondria (mitophagy) using the established OM45-GFP assay^31^. In this assay, the outer mitochondrial membrane (OMM) protein OM45 is tagged with a C-terminal GFP. Once OM45-GFP is delivered to the vacuole during mitophagy, OM45 is rapidly degraded resulting in the production of free GFP. We performed the OM45-GFP processing assay to assess the effect of the putative amphipathic helix deletion on mitophagy. While we observed a slight drop in the level of free GFP produced at 2 hours, this change was not significant suggesting that deletion of the amphipathic helix from Atg11 does not impair mitochondrial degradation by autophagy (**Figure 2D** and **E**). Furthermore, immunoblotting against 2xGFP-Atg11 and 2xGFP-Atg11^Δ612-646^, expressed from their endogenous promoters in *atg11Δ* cells, showed that the deletion of amino acids 612-646 did not alter the expression of Atg11, suggesting that deletion of this helix did not destabilize Atg11 (**Figure 2F**).

Atg32, an integral membrane protein of the OMM, serves as the selective autophagy receptor (SAR) for mitophagy in yeast^32,33^. Under mitophagy-inducing conditions, we and others have shown that Atg32 clusters on the mitochondrial surface, forming discrete punctum^27,34^. These Atg32 puncta initially colocalize with Atg11 and subsequently with the autophagosomal marker Atg8, before being transported to the vacuole^32,35^. Thus, these puncta are thought to represent sites of mitophagy initiation. In contrast to OM45-GFP, which shows only a faint vacuolar GFP signal after 2 hours of starvation, we and others have observed that Atg32 clustering into mitophagy initiation sites occurs more rapidly, with GFP-Atg32 puncta forming within 30 minutes of starvation^27,35^. This rapid clustering suggests that monitoring Atg32 puncta formation may provide a more effective approach for studying the early stages of mitophagy initiation. We therefore tested to see if Atg11^Δ612–646^ altered the formation of Atg32 puncta. To test if Atg11^Δ612–646^ altered the kinetics of mitophagy initiation, we monitored the formation of GFP-Atg32 puncta, in response to nitrogen-starvation, in real-time using the temperature- and flow rate- controlled FCS2 chamber (**Figure 2G**). We found that the rate of GFP-Atg32 puncta appearance was significantly lowered in cells expressing Atg11^Δ612–646^ than in cells expressing WT Atg11 (**Figure 2H, Supplemental Movie 1)**. At 28 minutes post starvation, 97% of cells expressing WT Atg11 showed at least one GFP-Atg32 punctum, whereas only 43% of cells expressing Atg11^Δ612–646^ displayed at least one GFP-Atg32 punctum at the same time point (**Supplemental Movie 2**). Moreover, 2xGFP-Atg11^Δ612–646^ was still recruited to the RFP-Atg32 puncta, similar to 2xGFP-Atg11^Δ612–646^, suggesting that deletion of the amphipathic helix did not affect recruitment of Atg11 during mitophagy (**Figure S5**). Together, our OM45-GFP processing and GFP-Atg32 clustering results suggest that the deletion of the predicted amphipathic helix in Atg11 causes a delay in the kinetics of mitophagy initiation but does not alter the bulk degradation of mitochondria.

### Atg11 forms clusters in solution

Formation of autophagy protein condensates have been proposed to enhance weak protein interactions, direct the formation of protein-complexes, and play a role in the overall organization of autophagy initiation sites^36,37^. We observed that GFP-Atg11 clustered on membrane tubes and SLBs which prompted us to investigate if Atg11 may cluster in solution or if the clustering of Atg11 is induced by membrane binding. GFP-Atg11 was purified in the presence of 300 mM NaCl as this was previously reported to allow for soluble full-length Atg11(**Figure S1A**)^9,18^. We monitored 1 µM GFP-Atg11 by confocal microscopy to determine if it formed puncta, which would indicate clustering of Atg11 (**Figure 3A**). We found that while GFP-Atg11 appeared largely diffuse at 300 mM NaCl, it formed puncta when the NaCl concentration was reduced to 150 mM or less (**Figure 3A and B**). We next asked if the formation of these puncta was dependent on the concentration of Atg11. We diluted GFP-Atg11 to concentrations from 0.05 µM to 5.0 µM into buffer with a final concentration of 150 mM NaCl (**Figure 3C**). We observed that increasing the concentration of Atg11 increased the number of Atg11 puncta indicating that Atg11 clustering is both salt and concentration dependent (**Figure 3D**). The puncta we observed were not liquid-like condensates as they are not entirely circular, nor did we observe the fusion of individual puncta. It is possible that diluting Atg11 into 150 mM NaCl resulted in an irreversible aggregation event. Therefore, to test if the formation of Atg11 clustering was reversible, we first generated Atg11 puncta in 150 mM NaCl, and then increased the NaCl concentration to 300 mM by mixing the 1 µM Atg11 in 150 mM NaCl with 1 µM of Atg11 in 450 mM NaCl (**Figure 3E**). After the samples were mixed, we saw a near complete loss in the formation of Atg11 puncta suggesting that these Atg11 clustering events are reversible in solution (**Figure 3F**). We next wanted to determine whether Atg11-Nterm or Atg11-Cterm could undergo similar clustering events as full-length Atg11 in solution. Therefore, we diluted GFP-Atg11, GFP-Atg11-Nterm and GFP-Atg11-Cterm into buffer containing 150 mM NaCl. While GFP-Atg11 formed puncta at 1 µM in 150 mM NaCl, neither the Nterm nor Cterm regions of Atg11 formed puncta at 1 µM in 150 mM NaCl, suggesting that the ability of Atg11 to cluster is a property of the full-length protein (**Figure 3G and H**). This data demonstrates that while full-length Atg11 can form reversible clusters in solution that are salt and concentration dependent, the Nterm and Cterm regions of Atg11 alone are unable to form clusters. This is consistent with our membrane binding observations in **Figure 1** where full-length Atg11 bound to membranes in a clustered fashion while the Nterm and Cterm regions of Atg11 bound membranes in a diffuse manner.

**Figure 3.**
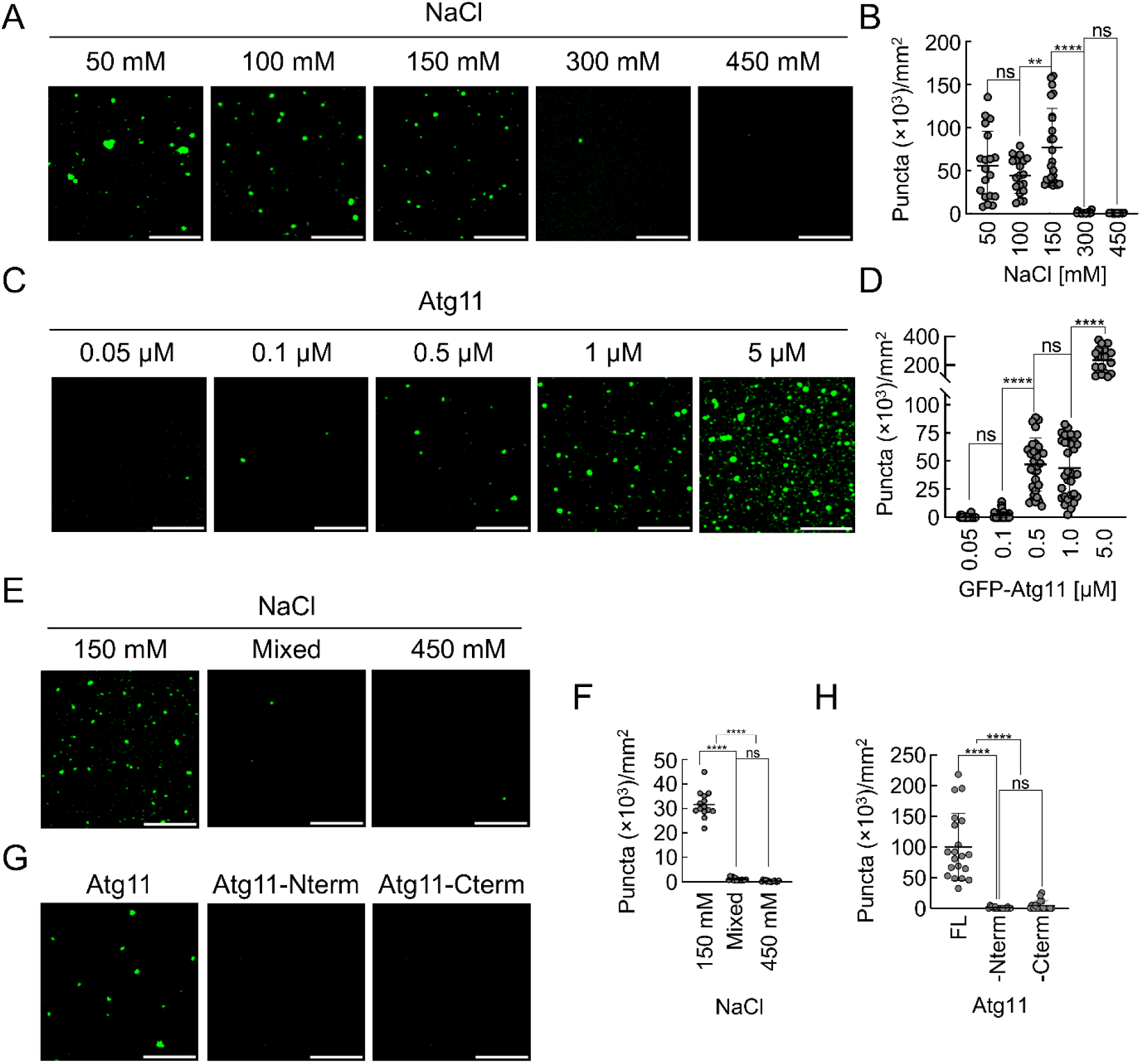
Atg11 self-associates in a salt and concentration dependent manner. Purified GFP-Atg11 was diluted into different buffers and imaged by confocal microscopy. (**A**) GFP-Atg11 puncta formation at 1uM protein concentration in different NaCl concentrations (as indicated). Scale bar=5μm. (**B**) Quantification of A. (**C**) GFP-Atg11 puncta formation in 150 mM NaCl at different protein concentrations (as indicated). Scale bar=5μm. (**D**) Quantification of C. (**E**) GFP-Atg11 was diluted to 1 μM in 150 mM or 450 mM NaCl, then imaged. Samples were mixed to produce 300 mM NaCl at 1 μM protein concentration. (**F**) Quantification of E. (**G**) Representative images of clustering observed when GFP-Atg11 constructs (FL, Atg11-Nterm or Atg11-Cterm) were diluted to 1 μM in 150 mM NaCl and imaged as before. (**H**) Quantification of clustering observed when GFP-Atg11 constructs (FL, Atg11-Nterm or Atg11-Cterm) were diluted to 1 μM in 150 mM NaCl and imaged as before. Each experiment was conducted in 3 independent repeats, with 7 images acquired at each repeat, amounting to a total area of 30492 μm^2^ per repeat. Puncta analyzed varied from ∼30 to ∼19,000, depending on the group. The number of puncta were normalized to the area imaged, yielding particles per mm^2^. The data are presented as means, with error bars indicating standard deviation. Groups were compared using a one-way ANOVA test with Tukey’s correction for multiple comparisons. * p< 0.05, ** p<0.01, *** p<0.001, **** p<0.0001, ns is not significant.

### The Atg32-Atg11 complex tethers autophagosomal-like liposomes to mitochondria-like GUVs

We have demonstrated that Atg11 can bind membranes in a clustered manner via a predicted amphipathic helix and that a loss of this helix leads to a delay in the formation of Atg32 puncta and the onset of mitophagy in yeast. Recruitment of the highly-curved Atg9 vesicles by the Atg32-Atg11 complex is a critical step in initiating autophagosomal membrane formation during mitophagy^7^. To investigate whether the ability of Atg11 to bind membranes enables the tethering of liposomes (mimicking the size and composition of Atg9 vesicles) to mitochondrial membranes, we utilized the <u>G</u>UV and <u>L</u>iposome <u>T</u>ethering (GLT) assay, which we recently developed^20^. GLT monitors tethering by measuring the co-localization of liposomes with GUVs in the presence and absence of potential tethering proteins. We made GUVs containing a lipid composition similar to the OMM (OMM-GUVs)^38–40^. GFP-Atg11 alone was unable to bind these OMM-GUVs further highlighting that Atg11 is not a broad membrane binding protein (**Figure S6A**). Next, we displayed the cytosolic domain of the mitophagy SAR Atg32 (1-381), which contains the Atg11 binding region, fused to an N-terminal GFP and a C-terminal 10xHistidine tag, on OMM-GUVs (**Figure S6B**)^41^. GUVs included 5 mol% of DGSNi^2+^NTA to recruit the cytosolic domain of GFP-Atg32_1-381_. To monitor if Atg11 can bind to these Atg32-displaying OMM-GUVs, we labeled purified Atg11 with Alexa Flour 594. Atg11 was recruited to GFP-Atg32_1-381_ bound GUVs. In contrast, no Atg11 recruitment occurred to GUVs displaying 6xHistidine tagged GFP (**Figure S6C)**, demonstrating that Atg11 requires Atg32 for its recruitment to the surface of OMM-GUVs (**Figure S6D**). To test if the Atg32-Atg11 complex could tether autophagosomal-like liposomes, we added sonicated automix+PI3P liposomes, mimicking the lipid composition and size of Atg9 vesicles (30 nm diameter), to GFP-Atg32_1-381_ OMM-GUVs in the presence and absence of Atg11 (**Figure 4A**). Automix+PI3P liposomes were recruited to Atg32_1-381_ bound GUVs only in the presence of Atg11 (**Figure 4B-E**). Fluorescence intensity traces along the GUVs further confirm that peaks of liposome fluorescence correspond with peaks of GFP-Atg32_1-381_ fluorescence and were only seen in presence of Atg11 (**Figure 4C** and **D**). Interestingly, tethered automix+PI3P liposomes appeared punctate on GUVs and these overlap with peaks of Atg32 fluorescence, suggesting that Atg11 may be able to cluster liposomes on the surface of cargo (**Figure 4C**). While Atg11 robustly tethered 30 nm automix+PI3P liposomes to Atg32 OMM-GUVs, the ability of Atg11 to recruit larger automix+PI3P liposomes (1 µm diameter) was significantly reduced (**Figure 4F** and **G**). This is in agreement with our results on SMrT tubes showing that automix+PI3P membrane binding by Atg11 occurred in a curvature-dependent manner.

**Figure 4.**
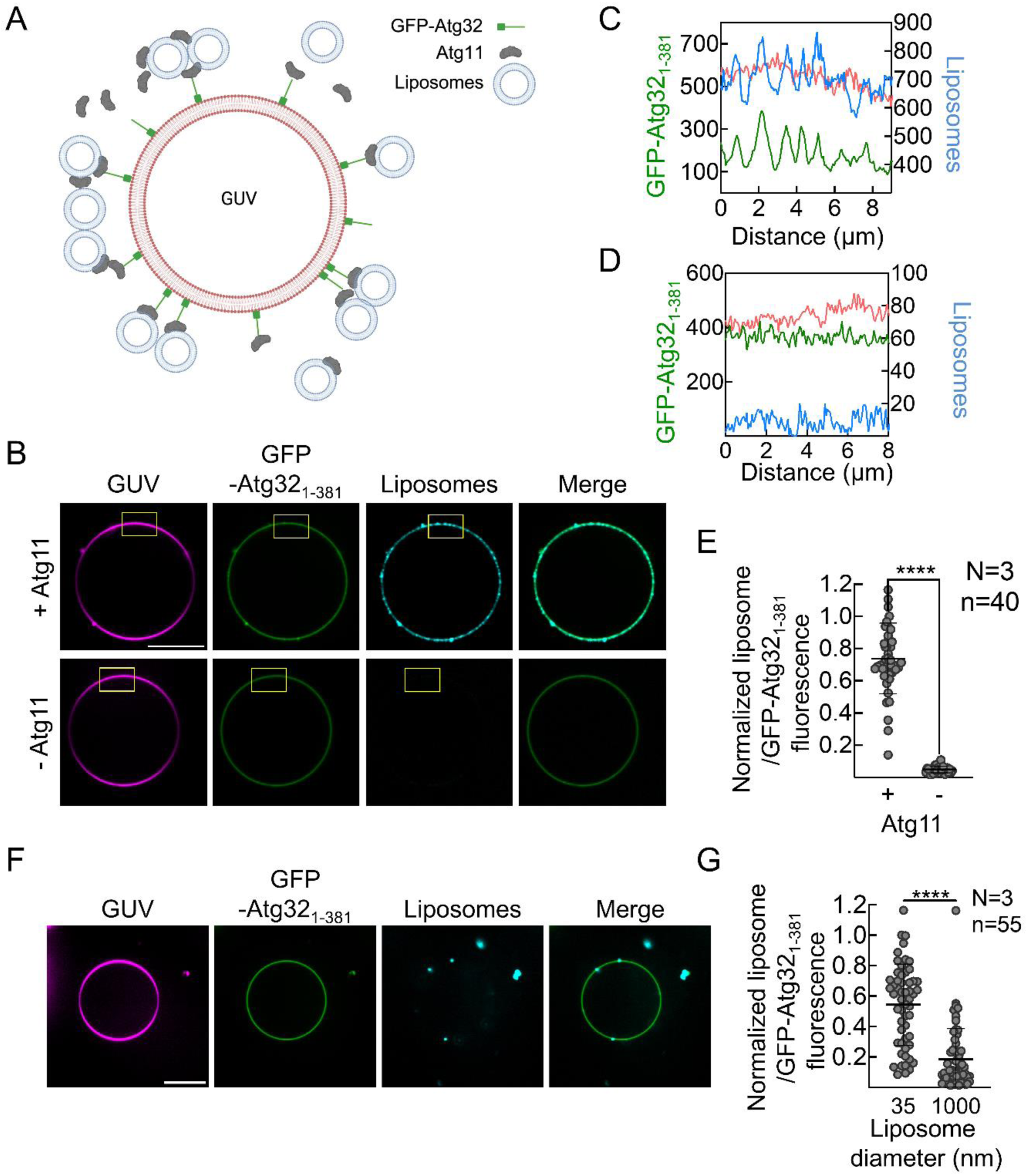
Atg11 and Atg32 coordinate to tether curved autophagosomal membranes to cargo mimetics. (**A**) Schematic showing tethering of autophagosomal membranes to mitochondrial mimetic GUV reconstituted using the GLT assay. OMM-GUVs containing mitochondrial lipids and DGS-Ni^+2^NTA (5 mol%) displaying the cytosolic domain of Atg32 (His-GFP-Atg32_1-381_) are used as a mitochondria mimic to recruit Atg11. 35nm-automix+PI3P liposomes are then added to reconstitute heterotypic tethering by the Atg32-Atg11 complex. (**B**) Representative fluorescence images of GLT with GUVs displaying His-GFP-Atg32_1-381_ which were sequentially incubated with Atg11 and automix+PI3P liposomes (top panel) or automix+PI3P liposomes alone (bottom panel). Scale bar=10μm. Yellow box marks the region used for tracing line profiles in C and D. (**C**) Fluorescence intensity profiles from a line drawn along the GUV in B (top panel) showing liposome and GFP-Atg32_1-381_ fluorescence intensities along the GUV. (**D**) Fluorescence intensity profiles from a line drawn along the GUV in B (bottom panel) showing liposome and GFP-Atg32_1-381_ fluorescence intensities along the GUV. (**E**) Quantification of the ratio of liposomes to GFP-Atg32_1-381_ fluorescence from B. Data are presented as means ± SD from 3 independent repeats (N=3). ****p<0.0001 analyzed using Mann-Whitney’s test. n indicates total number of GUVs analyzed per condition. (**F**) Representative image of a GUV displaying His-GFP-Atg32_1-381_ sequentially incubated with Atg11 and 1μm-automix+PI3P liposomes. (**G**) Quantification of the ratio of liposomes to GFP-Atg32_1-381_ fluorescence from F. Merge images show an overlay of GFP-Atg32_1-381_ and liposomes fluorescence channels. Data are presented as means ± SD from three independent repeats (N=3). n indicates the total number of GUVs analyzed per condition. **** indicates p<0.0001 analyzed using non-parametric Mann-Whitney’s test. OMM-GUVs were made with 69.9% DOPC, 10% DOPE, 10% DOPS, 5% cardiolipin, 5% DGS-Ni^+2^NTA, and 0.1% RhPE. Automix+PI3P were made of 43.9% PC, 19% PE, 32% PI, 5% PI3P with 0.1% DiD. Liposomes were either sonicated to obtain 30-50nm curved liposomes or extruded through a 1μm-filter.

We hypothesized that using YPL could override the need for curvature as Atg11 was able to bind GUVs containing YPL but not automix+PI3P GUVs (**Figure S2**). This may be due to the heterogeneity of lipids present in YPL which could lead to packing defects similar to what would be expected for more highly curved vesicles^48^. To test if Atg11 was also able to tether larger YPL liposomes to GUVs, we performed the GLT assay with 400 nm YPL liposomes and GFP-Atg32_1-381_ bound GUVs in the presence and absence of Atg11 (**Figure S7A**). As expected, YPL liposomes were tethered on GUVs only in the presence of Atg11 (**Figure S7D**) and the tethered liposomes colocalized with regions of higher Atg32 fluorescence (**Figure S7B** and **C**).

### The N-terminal region of Atg11 is critical for membrane tethering

Our results till now show that Atg11 binds membranes via a putative amphipathic helix located in its N-terminus and that it can self-associate to form clusters. Additionally, the interaction between Atg11 and Atg32 leads to membrane tethering in vitro. Therefore, to test if the Atg11-Nterm contributes to tethering of autophagosomal-like liposomes, we carried out GLT using GFP-Atg32_1-381_ bound OMM-GUVs and automix+PI3P liposomes with Atg11 (1µM) alone and Atg11(1µM) mixed with 20-fold molar excess of Atg11-Nterm (20µM) (**Figure 5A**). We reasoned that excess Atg11-Nterm would compete with the full-length Atg11 and inhibit tethering of automix+PI3P liposomes to GUVs. As a control we also carried out the GLT assay using Atg11 (1µM) mixed with 20-fold molar excess of GST (20µM). Quantification of the fluorescence intensity of liposomes normalized to that of GFP-Atg32_1-381_ showed that the addition of excess Atg11-Nterm led to a near complete block in automix+PI3P liposome tethering (**Figure 5B**). Atg11 mixed with an excess of GST had only a modest reduction in the tethering of automix+PI3P liposomes. This lends further support to the idea that autophagosomal-like liposome tethering by Atg11 occurs via membrane binding by the N-terminal region of Atg11.

**Figure 5.**
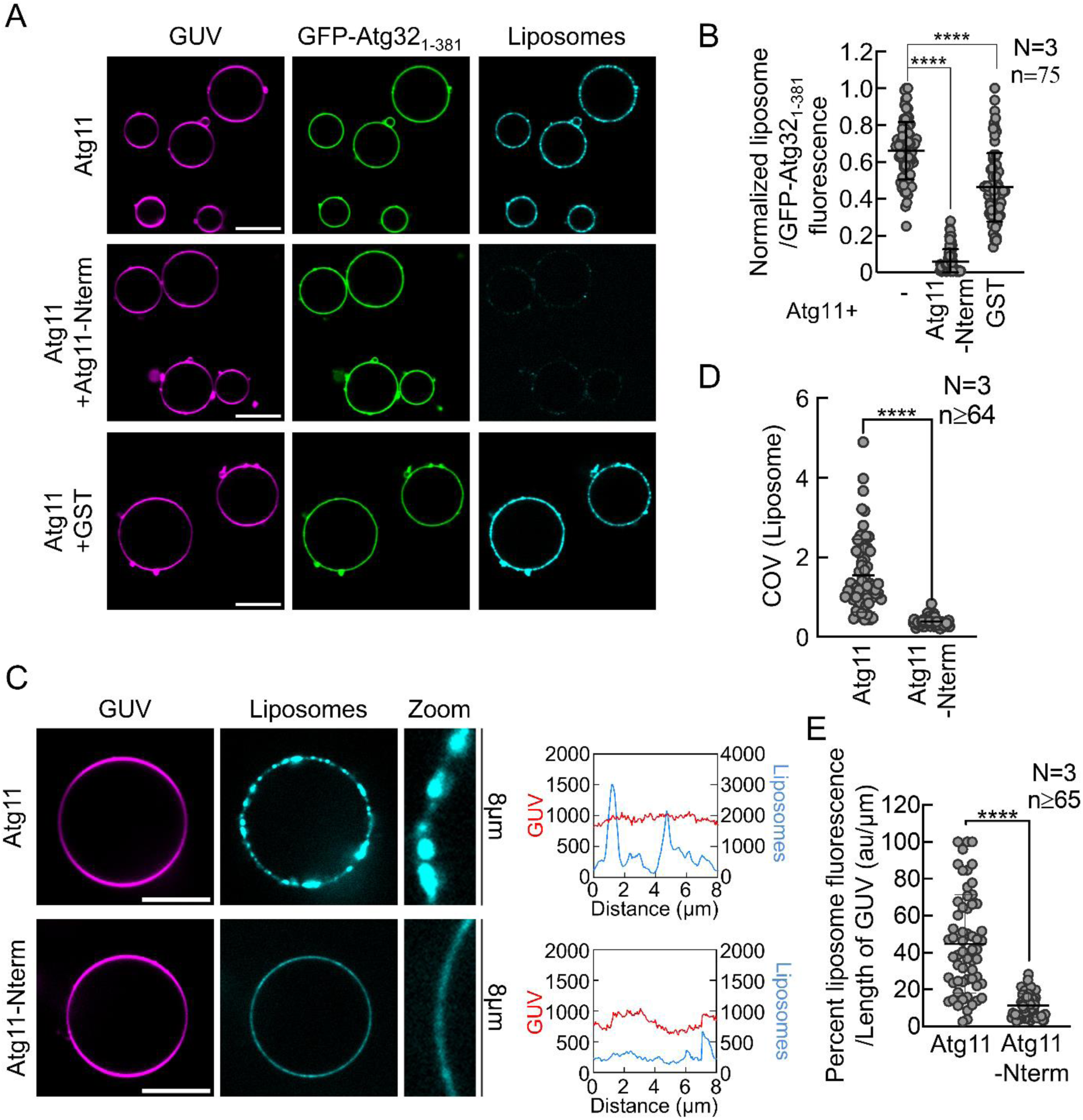
Membrane binding by Atg11-Nterm and self-association promotes autophagosomal vesicle tethering. OMM-GUVs displaying GFP-Atg32_1-381_ were incubated with Atg11 alone, Atg11 with 20-fold molar excess of Atg11-Nterm or GST and subsequently mixed with autophagosomal liposomes. (**A**) Representative images of autophagosomal liposome recruitment to OMM-GUVs with Atg11 alone, Atg11 with excess of Atg11-Nterm or Atg11 with excess of GST. Scale bar=10μm. (**B**) Quantification of the ratio of liposomes to GFP-Atg32_1-381_ fluorescence from A. Data from three independent experiments (N=3) are presented as mean ± SD. **** indicates p<0.0001 calculated using one-way ANOVA with Šídák’s multiple comparisons test. n indicates the total number of GUVs analyzed per condition. (**C**) Representative images of autophagosomal liposome recruitment to OMM-GUVs displaying GFP-Atg32_1-381_ with Atg11 or GFP-Atg11-Nterm alone (recruited via 6X-His tag). Scale bar=10μm. Fluorescence intensity profiles for the GUV and liposome channels along the circumference of GUVs from the zoomed inset are shown to the right of each representative image. (**D**) Quantification of the coefficient of variance (COV) for liposome fluorescence along the circumference of the GUV from. Data are presented as mean ± SD from three independent repeats (N=3). n indicates the total number of GUVs analyzed per condition. **** indicates p<0.0001 analyzed using the non-parametric Mann-Whitney’s test (**E**) Quantification of the ratio of liposomes to GFP-Atg32_1-381_ fluorescence from C. Data are presented as mean ± SD from three independent repeats (N=3). n indicates the total number of GUVs analyzed per condition. **** indicates p<0.0001 analyzed using non-parametric Mann-Whitney’s test. OMM-GUVs were made with 69.9% DOPC, 10% DOPE, 10% DOPS, 5% cardiolipin, 5% DGS-Ni^+2^NTA, and 0.1% RhPE. Sonicated Automix+PI3P were made of 43.9% PC, 19% PE, 32% PI, 5% PI3P with 0.1% DiD.

Next, we compared the liposome tethering abilities of full-length Atg11 with that of the Atg11-Nterm. Full-length Atg11 was mixed with GFP-Atg32_1-381_ bound OMM-GUVs and subsequently incubated with automix+PI3P liposomes (**Figure 5C**, top panel). Because the Atg32 binding interface lies at the C-terminus of Atg11, we directly recruited Atg11-Nterm to NTA-containing OMM-GUVs and subsequently added automix+PI3P liposomes. GUV-bound Atg11-Nterm was also able to tether liposomes further supporting the role of the Atg11-Nterm in membrane binding and tethering (**Figure 5C**, bottom panel). Intriguingly, while liposomes tethered via full-length Atg11 showed a punctate distribution, liposomes tethered via Atg11-Nterm uniformly coated the GUVs (**Figure 5C** zoom images). Fluorescence intensity traces along the GUVs revealed that automix+PI3P liposomes tethered by full-length Atg11 showed discreet peaks. In comparison, liposomes tethered via Atg11-Nterm had a lower and more uniformly distributed intensity (**Figure 5C**). To quantitate the distribution of liposomes, we compared the coefficient of variance (COV) of liposome fluorescence, calculated as standard deviation by mean, which serves a measure for the extent of liposomes clustering. We found that the COV for Atg11-Nterm is significantly lower than the COV for full-length Atg11 (**Figure 5D)**, suggesting that while OMM-GUV bound Atg11-Nterm alone can tether autophagosomal-like liposomes, it fails to cluster these liposomes. Importantly, the liposome fluorescence per micron calculated along the surface of GUVs was significantly higher for full-length Atg11 than Atg11-Nterm, suggesting that the clustering by Atg11 enhances the tethering of autophagosomal-like liposomes (**Figure 5E**).

## DISCUSSION

Our results demonstrate, for the first time, that Atg11 can bind highly curved membranes containing an autophagosomal-like lipid composition. Intriguingly, we observed that Atg11 membrane binding occurred in a punctate manner and that this clustering of Atg11 could be recapitulated in the absence of membrane. We also identified a putative amphipathic helix in the N-terminal region of Atg11 as the primary membrane binding interface. We show that the deletion of this helix results in a delay in the formation of mitophagy initiation sites in yeast by monitoring the formation of Atg32 puncta in real-time. To further explore the role of Atg11 membrane binding, we used a method we recently developed—GLT—to reconstitute membrane tethering in vitro^20^. Using GLT, we demonstrated that Atg11, in complex with Atg32, recruits autophagosomal-like liposomes to OMM-GUVs. Comparing the tethering properties of Atg11 full-length and Atg11-Nterm, we show that the ability of Atg11 to cluster enhances liposome tethering and causes clustering of tethered liposomes on OMM-GUVs. Taken together our work suggests a possible role for Atg11 in the recruitment or organization of Atg9 vesicles during selective autophagy.

The interaction between Atg9 and Atg11 was previously shown to be only partially required for the recruitment of Atg9 vesicles to cargo in yeast^8^. Our results showing that Atg11 can bind autophagosomal-like membranes directly and tether them to OMM-GUVs, could provide a mechanistic explanation to this observation. Given these results it is possible that mitochondrial-recruited Atg11 may tether Atg9 vesicles to mitochondria through dual interactions—binding both Atg9 and autophagosomal membranes. While the deletion of the membrane binding helix of Atg11 results in a delay in the formation of Atg32 puncta, this deletion did not significantly hamper delivery to the mitochondrial protein OM45 to the vacuole at 2 hours post mitophagy induction. Therefore, we propose that Atg11 membrane binding via its amphipathic helix is critical for the initial recruitment of Atg9 vesicles during the early stages of mitophagy (**Figure 6**). Once the low-affinity protein interactions at mitophagy initiation sites are stabilized via avidity, membrane binding via the amphipathic helix may become less significant. During this initial phase, Atg11 membrane association via the amphipathic helix may: 1) work synergistically with Atg11-Atg9 interaction to enhance or organize Atg9 vesicle recruitment, or 2) act as co-incident targeting mechanism ensuring selective binding of Atg9 vesicles specifically at autophagy initiation sites. However, which of these possibilities is correct will need to be further explored in subsequent work.

**Figure 6.**
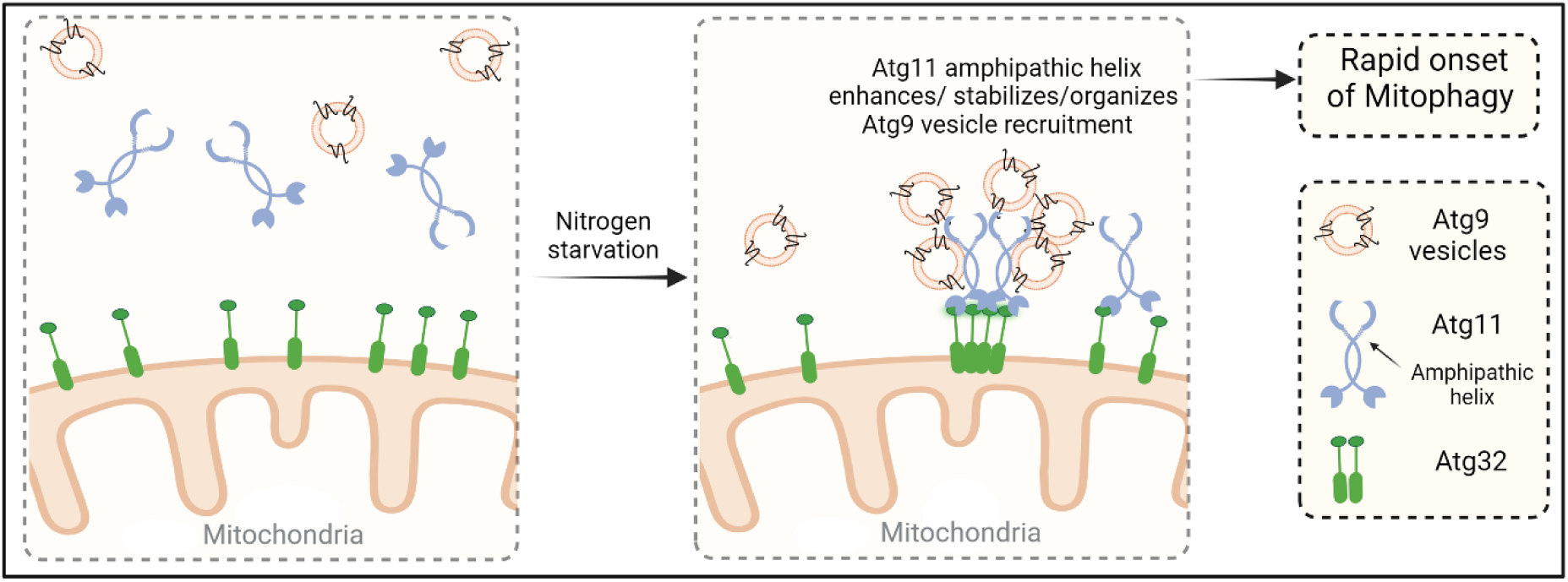
Proposed role of Atg11 membrane binding in mitophagy initiation. In response to nitrogen starvation, Atg32 clusters and recruits Atg11 on mitochondria. An amphipathic helix in Atg11 engages with negatively charged lipids in Atg9 vesicles. This membrane binding may serve to enhance, stabilize or organize the recruitment of Atg9 vesicles helping in rapid onset of mitophagy.

We also demonstrate that GLT can be used for reconstituting the earliest steps of organelle selective autophagy. We propose that GLT is a useful alternative to other reconstitution methods that rely on selective autophagy receptors immobilized on Sepharose or agarose beads which may more accurately mimic protein-based cargos such as aggregated proteins^9,42^. In contrast to Sepharose or agarose-based reconstitution methods, selective autophagy receptors immobilized on GUVs can diffuse freely along the bilayer and provide a more biologically relevant system for studying organelle selective autophagy initiation where integral membrane selective autophagy receptors such as Atg32 can diffuse along the cargo membrane.

Our work demonstrating that Atg11 can directly interact with membranes contrasts a previous study reporting that Atg11 does not bind membranes^7^. However, two key differences could account for the observed discrepancy in the membrane binding abilities of Atg11. First, we purified Atg11 using Freestyle 293 cells as a recombinant expression system where Atg11 appears to be a dimer similarly to SF9 insect cells, while the Matscheko et al. study used monomeric Atg11 purified from *E. coli* (**Figure S1B**). Second, we used liposomes containing YPL and autophagosomal-like lipid compositions. Both mixtures contain approximately 42% negatively charged lipids. In contrast, monomeric Atg11 was previously shown to be unable to bind liposomes composed of 60% POPC, 20% Cholesterol, 10% POPE and 10% POPS which contain only 10% negatively charged lipids^7^. We have previously demonstrated that Atg13, a member of the autophagy initiation machinery, binds liposomes containing at least 42.5% negatively charged lipids^20,43^. These findings may also suggest a common mechanism among autophagy proteins, where membrane association occurs preferentially with the highly negatively charged autophagosomal membranes, rather than the less charged membrane of the cargo. This differential lipid composition may facilitate the spatial organization of autophagy proteins during autophagosome expansion.

Our data shows that the deletion of the amphipathic helix in Atg11 reduces but does not completely abolish membrane binding. Interestingly, the mammalian homolog of Atg11, FIP200, has been shown to interact with membranes via its large coiled-coil (CC) domain^17^. Our experiments with SMrT tubes show a weak membrane binding for the Atg11 C-terminus, raising the possibility that Atg11 has additional weaker membrane binding regions located in its C-terminus. Given the enhanced recruitment and clustering of liposomes observed in the presence of full-length Atg11 but not Atg11-Nterm it is possible that secondary membrane binding sites might further strengthen Atg11 membrane binding and clustering.

While this work was being completed, a recent study demonstrated that the diffusion of selective autophagy receptors on the cargo surface promotes the recruitment and phase separation of Atg11 into liquid-like droplets. These phase-separated assemblies, stabilized by dynamic weak interactions, are proposed to stabilize the initiation machinery through avidity while providing spatio-temporal regulation to phagophore initiation^44^. Our findings on Atg11 clustering on autophagosomal membranes, and the enhancement of autophagosomal-like liposome tethering to OMM-GUVs by Atg11 clustering, are in close agreement with this model. Intriguingly, the properties of Atg11 clustering that we observed appear to be similar to those reported in Licheva et al. in that Atg11 began to form significant clusters at 0.5 µM in 150 mM NaCl (**Figure 3**). However, Atg11 clusters we observed are much smaller, less circular and do not appear to be liquid-like as was described in Licheva et al. This may be due to differences in the expression systems used or the imaging conditions. We purified Atg11 using the Freestyle 293 expression system, whereas Licheva et al. purified Atg11 from SF9 insect cells. Additionally, our imaging was performed on glass coverslips, while Licheva et al. used a six-channel chamber. Despite these differences both our study and the Licheva et al. study point towards the importance of Atg11 higher order assemblies (either clustering or phase separation) in the initiation of selection autophagy.

There is now a significant body of work that highlights the clustering of selective autophagy receptors into high order assemblies to facilitate selective autophagy initiation^5^. In addition, several studies also demonstrate a critical role for higher order assembly, primarily phase separation, for the assembly of the autophagy initiation machinery^37,44^. Our work extends this understanding by demonstrating that Atg11 can form clustered scaffolds on membranes. The clustering of Atg11 also enhances the recruitment of and organization of small autophagosomal-like vesicles on membrane-based cargo mimetics. Therefore, it is plausible that weak multivalent interactions of Atg11 with the membrane, with itself and other autophagy proteins including selective autophagy receptors may all contribute to the recruitment and organization of membrane during selective autophagy initiation.

## MATERIALS AND METHODS

### Recombinant protein expression and purification

All constructs used for protein purification in this paper are listed in **Table S1**. StrepII-GFP-Atg32(1-381)-10XHis were recombinantly expressed in *E. coli* Star cells (Invitrogen C601003). Cultures were grown in LB broth (Fisher Scientific, BP1426) to an OD600 of 0.6 to 0.8 at 37°C while shaking at 210 rpm. Cultures were cooled at 4°C for 20 min. Protein expression was induced by adding 1 mM isopropylthio-β-D-galactopyranoside (IBI Scientific, IB02125) and the cultures were grown for an additional 18 h at 18°C. Cells were harvested and cell pellets were stored at −80°C. For purification of StrepII-GFP-Atg32(1-381)-10XHis, cell pellets were thawed and resuspended in 50 mM Tris pH 8.0, 500 mM NaCl, 2.5mM imidazole, 5 mM MgCl_2,_ 1 mM phenylmethanesulfonyl fluoride (PMSF; VWR, 0754) and cOmplete Mini EDTA-free protease inhibitor tablets (Roche, 11836170001). Cells were lysed using 3 strokes of a microfluidizer at 18, 000 psi. Triton X-100 (VWR, AAA16046-AE), was subsequently added to lysates at 1% vol/vol. Lysates were spun at 25,000 g for 30 minutes. The supernatant was then applied to pre-equilibrated Talon resin (Clontech, 635504). The resin was washed with 50 mM Tris pH 8.0, 500 mM NaCl, 10 mM imidazole. Bound proteins were eluted with 50 mM Tris pH8.0, 500 mM NaCl, 250 mM imidazole. Elutions from Talon resin were pooled and incubated with pre-equilibrated Strep resin (IBA, 2-5030-025). The resin was washed with 20 mM Tris pH 8.0 and 300 mM NaCl. Protein was eluted with 20 mM Tris pH 8.0, 300 mM NaCl and 50 mM biotin. Peak fractions containing protein were dialyzed against 20 mM Tris pH 8.0, 300 mM NaCl and 0.2 mM TCEP. Dialyzed protein was aliquoted, flash-frozen in liquid nitrogen and stored at −80°C until use.

Atg11 and GFP-Atg11, both with an N-terminal 2XStrep II tag, were recombinantly expressed in Freestyle cells for 3 days post transfection. Cells were harvested by spinning by 1000 g for 15 min. Cell pellets were flash frozen in liquid nitrogen and stored at −80°C.

Cell pellets were resuspended and thawed in 50 mM Tris pH 8.0, 500 mM NaCl, 1 mM DTT, 1 mM PMSF and a cOmplete Mini protease inhibitor tablet (Roche, 11836170001). Cells were lysed using 3 strokes of a microfluidizer at 10, 000 psi. Triton X-100 (VWR, AAA16046-AE), was subsequently added to lysates at 1% vol/vol. Lysates were spun at 25,000 g for 30 min. Supernatant was applied to Strep resin (IBA, 2-5030-025) at 1 mL/min using a peristaltic pump. Resin was washed with 20 mM Tris pH 8.0, 300 mM NaCl and 1 mM DTT. Bound protein was eluted with 20 mM Tris pH 8.0, 300 mM NaCl, 50 mM biotin and 1 mM DTT. Fractions containing protein were pooled and concentrated to 2 μM and flash-frozen in liquid nitrogen. For GFP-Atg32 binding experiments, Atg11 was fluorescently labeled with a three-fold molar excess of Alexa Fluor™ 594 C5 Maleimide (Invitrogen, A10256) for 1 h at room temperature. Labelling was quenched using 1 mM DTT. Unreacted dye was removed by dialyzing the labelled protein against 20 mM Tris pH 8.0, 300 mM NaCl, 0.2 mM TCEP and 1 mM DTT for 48 h. Fresh buffer was added after 24 h of dialysis.

### Yeast strains, plasmids and culture conditions

Δ*atg11*cell line was obtained from the *S. cerevisiae* knockout collection (Invitrogen). TKYM22 (SEY6210 OM45-GFP::TRP1) was a gift from Daniel Klionsky^45^. TKYM22 atg11Δ::NAT (XXY003) was generated by replacing atg11 with the NAT cassette from pAG25 in TKYM22^27^. GFP-Atg32 and RFP-Atg32 (pRS415-MET25) plasmids were kind gifts from Dr. Benedikt Westermann^46^. Plasmid expressing 2XGFP-Atg11 (yCPLAC33) and Atg11 (yCPLAC33) were reported earlier^27^. Atg11^Δ612–646^ was generated from Atg11 (yCPLAC33) by site directed mutagenesis. Cells were grown in SMD (0.67% yeast nitrogen base, 2% glucose and supplemented with the appropriate amino acid and vitamin mixture).

### Live imaging of GFP-Atg32 puncta formation

For GFP-Atg32 puncta formation experiments, cells expressing GFP-Atg32 under the MET25 promoter were grown to mid-log phase in SMD supplemented with 10 mg/mL methionine. GFP-Atg32 expression was then induced by shifting cells to SMD without methionine for 1hr. Subsequently, cells were transferred onto a concavalinA-coated glass coverslip assembled in a temperature- and flow-controlled FCS2 chamber (Bioptechs). Chamber temperature was maintained at 30 °C. To induce mitophagy, SD-N (0.17% yeast nitrogen base without amino acids and ammonium sulfate; 2% glucose) prewarmed to 30 °C was flowed into the chamber at a constant flowrate. Chamber was equilibrated with SD-N for 3 mins. Subsequently, timelapse images were acquired at every 4 mins interval for a total of 30-40 mins.

### OM45-GFP processing assay

The OM45-GFP processing assay was performed as described previously^31^. XXY003 cells were transformed with Atg11 and Atg11^Δ612–646^. Cells were grown at 30 °C in SMD to an OD600 of 0.8– 1.0. Cells were pelleted at 1000*g* for 10 min, washed and resuspended in SML (0.67% yeast nitrogen base, 2% lactic acid and supplemented with the appropriate amino acid and vitamin mixture) and grown for 16 h. Subsequently, cells were pelleted and resuspended in SD-N for 6 h. Finally, cells were harvested, lysed by bead beating and GFP was detected by western blot.

### Liposome preparation

All lipids were purchased from Avanti Polar lipids. Chloroform stocks of lipid mixtures at a desired ratio containing 0.1 mol% of the fluorescent lipophilic dye DiD were aliquoted in a clean glass tube and dried under a gentle nitrogen stream until all the chloroform had evaporated. Residual chloroform was removed under vacuum for 18 h. Dried lipids were resuspended in 20 mM Tris pH 8.0, 100 mM NaCl to a final concentration on 1 mM. Automix+PI3P liposomes were sonicated using a probe sonicator at 10% amplitude for 5 mins with 1 sec on and 2 sec off. Sonication was carried out in an ice-water bath to prevent heating. 1-μm liposomes were prepared using an Avanti Mini Extruder using 1-μm Nuclepore Track-Etched membrane filters (Sigma Aldrich, WHA10417104).

### Giant unilamellar vesicle preparation

GUVs were made using the gel-assisted formation method^51^. Briefly, chloroform stocks containing lipids mixed at desired ratios containing the fluorescent lipid RhPE were aliquoted in a glass tube. Chloroform was dried under a stream of nitrogen followed by drying under vacuum (1 h). The dried lipids were resuspended in fresh chloroform to make a 1 mM lipid mixture. Polyvinyl alcohol (PVA, Mw = 14500; Sigma, 814894) was dissolved in boiling water to make a 5% stock and degassed in a vacuum chamber. PVA solution (10 μL) was spread into a thin film on a glass coverslip kept on a heating block set to 55°C. Chloroform-dissolved lipid mix (10 μL) was then spread on the dried PVA film. The dried PVA-lipid films were peeled from the glass coverslips and transferred to a 1.5-mL microcentrifuge tube. Films were hydrated in 300 μL assay buffer (20 mM Tris pH 8, 100 mM NaCl) for 20 min, and GUVs were released by gentle tapping. The PVA films were removed using a pipette.

### GUV liposome tethering (GLT)

For GLT experiments, GUVs (120 to150 μL) were first mixed with tethering protein or control protein in a 1.5 mL microcentrifuge tube for 10 min at room temperature. GUV mixtures were first tapped to resuspend settled GUVs and subsequently incubated with 10 μL of liposomes (1mM) for 5mins at room temperature. Liposomes were used within 3 days of extrusion or sonication. The GLT mixtures (300 μL total volume) were transferred to a BSA-passivated LabTek chamber (Nunc, Z734853) for imaging. GUVs were allowed to settle for 5 min prior to imaging. LabTek chambers were treated with 3 M NaOH for 10 min followed by extensive washes with milliQ water. NaOH-treated chambers were then passivated by incubation with BSA (3 mg/mL). BSA was removed by three washes with the assay buffer. GLT experiments with Atg11 were performed in 20 mM Tris pH 8, 100 mM NaCl. All proteins were spun at 100,000 g prior to use to remove any aggregates.

### Atg11 binding on SMrT tubes

SMrTs were prepared as described earlier^22,23^. Briefly, 3 μL of chloroform-dissolved automix and automix+PI3P stocks (1mM) were spread on a PEGylated coverslip. Chloroform was dried using a gentle stream of nitrogen. A ∼35 μL flow cell (Bioptechs) was assembled by placing a 0.1 mm silicone spacer between the PEGylated coverslip and an ITO-coated slide. Rehydration of the dried lipid film with 20mM Tris pH 8.0 and 100mM NaCl yielded supported lipid bilayers and vesicles. Lipid nanotubes were extruded from the vesicles using buffer flow. Subsequently, 300 μL of GFP-Atg11 (0.5 μM), GFP-Atg11-Nterm (2 μM) or GFP-Atg11-Cterm (2 μM) were incubated with SMrTs for 10 mins. Excess protein was washed with 3×300 μL of buffer (20mM Tris pH 8 and 100mM NaCl) and SMrTs were imaged. All proteins were spun at 100,000 g prior to use to remove any aggregates.

### Atg11 Clustering Assay

All buffers used for these assays contained 20 mM Tris pH 8.0 at 4°C and the indicated NaCl concentrations. Purified GFP-Atg11 was thawed on ice and centrifuged at 20,000 x g for 15 minutes at 4°C. GFP-Atg11 was diluted to a concentration of 1 µM in buffer containing 50, 100, 150, 300 or 450 mM NaCl, or to a concentration of 0.05, 0.1, 0.5, 1 or 5 µM in buffer containing 150 mM NaCl. After dilution 40 µl of sample was spotted onto a glass coverslip, mounted on a 63 X oil immersion objective on a Zeiss LSM 880 microscope. Samples were left on the microscope for 5 minutes before imaging. All images were acquired within 10 minutes after the sample was applied to the coverslip to prevent evaporation and thus concentration of the sample. For reversibility assays, GFP-Atg11 was diluted to 1µM in buffer containing 150 mM NaCl or 450 mM NaCl and imaged as described above. The remaining samples of 1µM GFP-Atg11 in 150 mM NaCl and 450 mM NaCl were mixed in a 1:1 ratio to generate 1µM GFP-Atg11 in 300 mM NaCl. For the comparison of GFP-Atg11, GFP-Atg11-Nterm and GFP-Atg11-Cterm protein was diluted to 1 µM in buffer containing 150 mM NaCl and imaged as described above.

### Fluorescence microscopy

Atg11 clustering was imaged on a Zeiss Airyscan LSM 880 microscope using 63 X oil immersion objective using Zeiss Immersol 518 F immersion oil at room temperature. GFP was excited using a 488 argon laser. Emission was detected using a Zeiss GaAsP-PMT detector.

All other images were acquired using a Nikon TiE microscope fitted with a Yokogawa CSU-W1 spinning disk system using a 100 X, 1.45 NA oil immersion objective. Images were acquired with a Photometrics Prime BSI sCMOS camera using the NIS elements software.

### Image analysis

Fluorescence microscopy images were analyzed using Fiji^52^.

To analyze tethering in GLT, images were first background corrected by subtracting the mode intensity value. Percent liposome fluorescence per length of GUV was calculated by normalizing the liposome fluorescence, collected from a two-pixel wide segmented line drawn along a GUV, to the length of the GUV. For calculating the liposome/protein fluorescence ratio, fluorescence intensities in liposome and protein channels from a two-pixel wide segmented line drawn along a GUV were collected. Mean liposome fluorescence was divided by mean protein fluorescence to yield liposome/protein fluorescence ratios.

For analysis of protein bound on SMrTs, images were background corrected by subtracting the mode value. Integrated density of protein and membrane fluorescence was measured from a 20×10 pixel rectangle placed centrally on the tubes or from a 2 to 4 μm^2^ square box placed on SLBs.

GFP-Atg32 puncta were analyzed by placing a segmented line centered on the punctum. Only those GFP-Atg32 puncta that were over 2.5-fold intensity of the background mitochondrial fluorescence intensity were counted.

For Atg11 clustering analysis images were analyzed in Fiji. A threshold was set for each image based on the average background intensity of the image. The analyze particle command was used with parameters of 0.2 µm to infinity for the particle size and 0.8-1.0 for the circularity.

### Statistical analysis

Statistical analysis was performed using GraphPad Prism (version 5.0a).

All experiments were performed in independent triplicates or duplicates. Liposomes and GUVs were prepared independently for each GLT experiment using freshly aliquoted lipids. Data from all repeats were pooled and plotted as scatterplots. Error bars represent standard deviation. Significance was calculated using the non-parametric Mann-Whitney’s test, one-way ANOVA with Tukey’s correction for multiple comparisons or Šídák’s multiple comparisons test, or t test as appropriate.

## Author Contributions

D.A., S.K., K.M.B., and M.J.R. conceived and planned the experiments. D.A., S.K., S.I.N. and K.M.B. carried out the experiments. D.A. and M.J.R. drafted the manuscript. All authors edited the manuscript.

## Acknowledgements

This work was supported by the National Institute of General Medical Sciences (NIGMS) grant R35GM128663 to M.J.R. Expression of Atg11 was performed in the BioMT service center and protein purification was performed in part performed at the BioMT Molecular Tools Core which are both supported by NIGMS grant P20GM113132. Sequencing of plasmids was performed by the Molecular Biology Shared Resource Center which is supported by NCI Cancer Center Support Grant P30CA023108. We would also like to thank Ann Lavanway for her work and support at the Dartmouth Imaging Facility. We thank Dr. Benedikt Westermann for the kind gift of GFP-Atg32 plasmid.

## Disclosure Statement

The authors have no conflicts of interest to declare.

## ABBREVIATIONS

Atg11: autophagy related 11
Atg32: autophagy related 32
DLS: dynamic light scattering
GLT: GUV liposome tethering assay
GUV: giant unilamellar vesicle
PE: phosphatidyl ethanolamine
PI3P: phosphatidyl inositol 3-phosphate
SAR: selective autophagy receptor
YPL: yeast polar lipids

## Notes

### Competing Interest Statement

The authors have declared no competing interest.

### Summary of Updates

In this most recent version figure 4 was removed based on the suggestion from reviewers that the manuscript should focus exclusively on the mechanisms of selective autophagy and not assay validation. The discussion has also been updated to acknowledge a recent manuscript which was published.

