## Supplemental Material for "Reconstitution of autophagosomal membrane tethering reveals that Atg11 can bind and cluster vesicles on cargo mimetics"

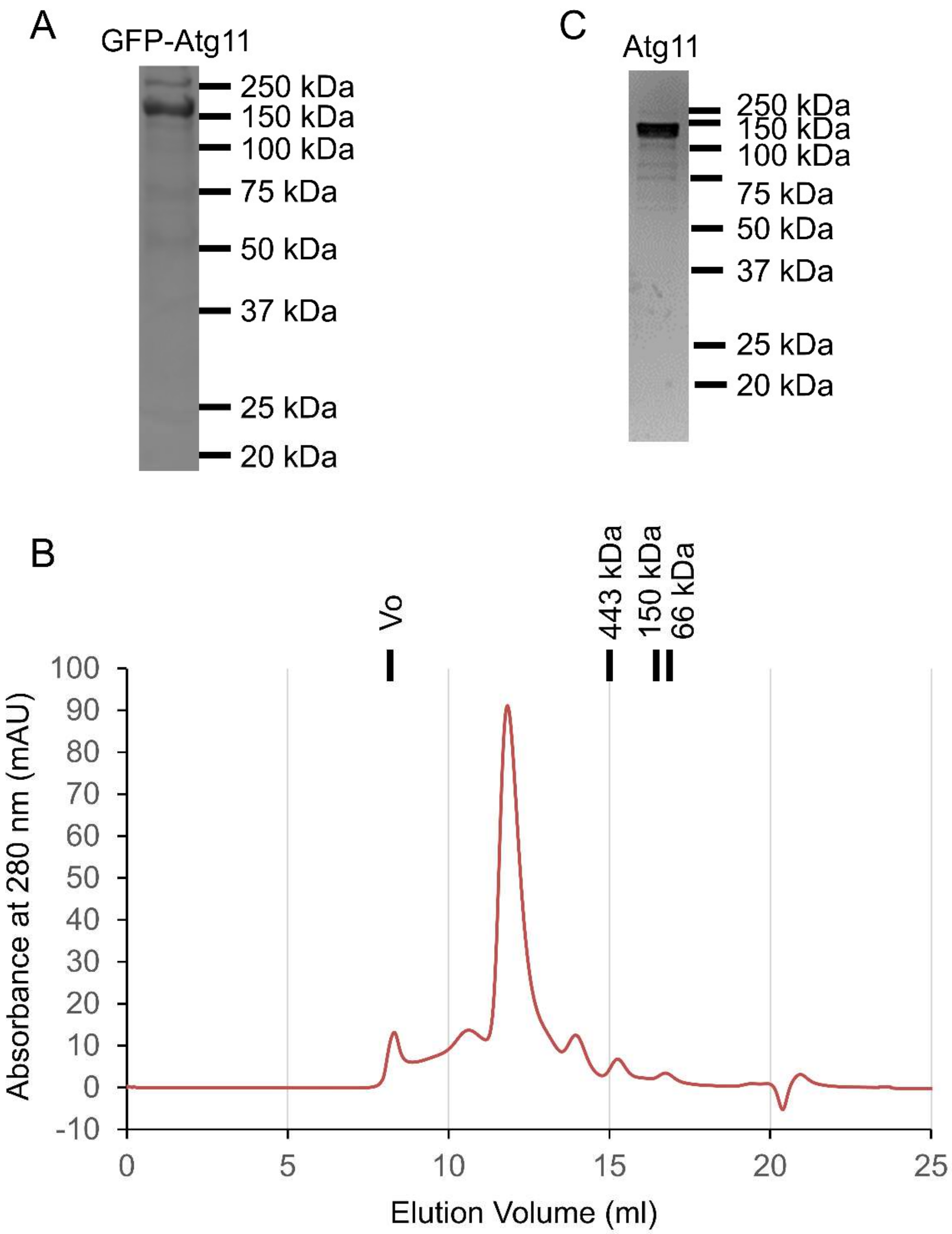


**Supplementary Figure 1. Purification and Characterization of Atg11 from Freestyle 293 Cells.** (**A**) Coomassie stained SDS-PAGE gel of purified Twinstrep-GFP-Atg11 from Freestyle293 cells. The monomer molecular weight of Twinstrep-GFP-Atg11 is 166.1 kDa. (**B**) SEC chromatography of Twinstrep-Atg11 purified from Freestyle 293 cells on a Superose 6 Increase 10/300 GL column in 20 mM Tris pH 8.0, 300 mM NaCl and 0.2 mM TCEP. The molecular weight of standards run on the same column in the same buffer are marked above the chromatograms. (**C**) Coomassie stained SDS-PAGE gel of Twinstrep-Atg11 from the center of the peak in the chromatogram in B. The monomer molecular weight of Twinstrep-Atg11 is 139.2 kDa.

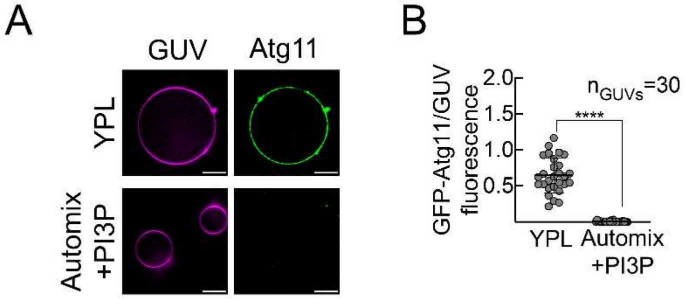


**Supplementary Figure 2. Atg11 binds YPL GUVs**. (**A)** Representative images showing binding of full-length GFP-Atg11 on GUVs containing yeast polar lipids (YPL, top panel) and autophagosomal-mimicking membrane with PI3P (automix+PI3P) (bottom panel). Scale bar=10µm. (**B**) Quantification showing mean±SD of GFP-Atg11 fluorescence normalized to membrane fluorescence from A. A total of 30 YPL and automix+PI3P GUVs each were analyzed (n=30). **** indicates p<0.0001 calculated using the non-parametric Mann-Whitney's test.

A

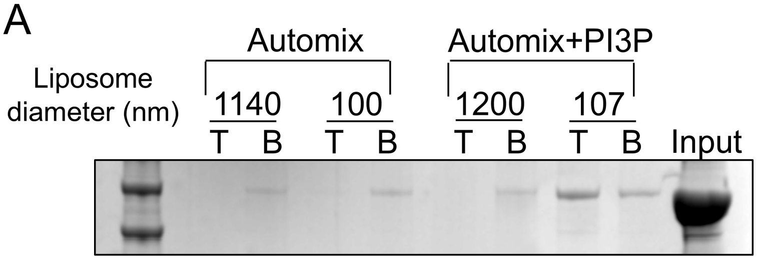


B

| | Liposomes | Liposomes+Atg11 |
|---|---|---|
| Diameter (nm) | 107 (+/- 15) | 260 (+/- 27) |

**Supplementary Figure 3. Validation of Atg11 membrane binding.** (**A**) Floatation assay with Atg11 and liposomes mimicking autophagosomal membranes with PI3P (automix+PI3P) and without PI3P (automix-PI3P). Liposome diameters indicate the sizes of liposomes before addition of Atg11. Top (T) and bottom (B) layers are shown in the representative SDS-PAGE gel. Input indicates total Atg11 used in floatation assay. (**B**) Sizes of automix liposomes with PI3P before and after addition of Atg11. Autophagosomal-mimicking membranes with PI3P (Automix+PI3P) were made of 43.9mol% PC, 19mol% PE, 32mol% PI, 5mol% PI3P with 0.1mol% RhPE. Autophagosomal-mimicking membranes without PI3P (Automix-PI3P) were made similarly except for the omission of PI3P and increase of PI to 37mol%.

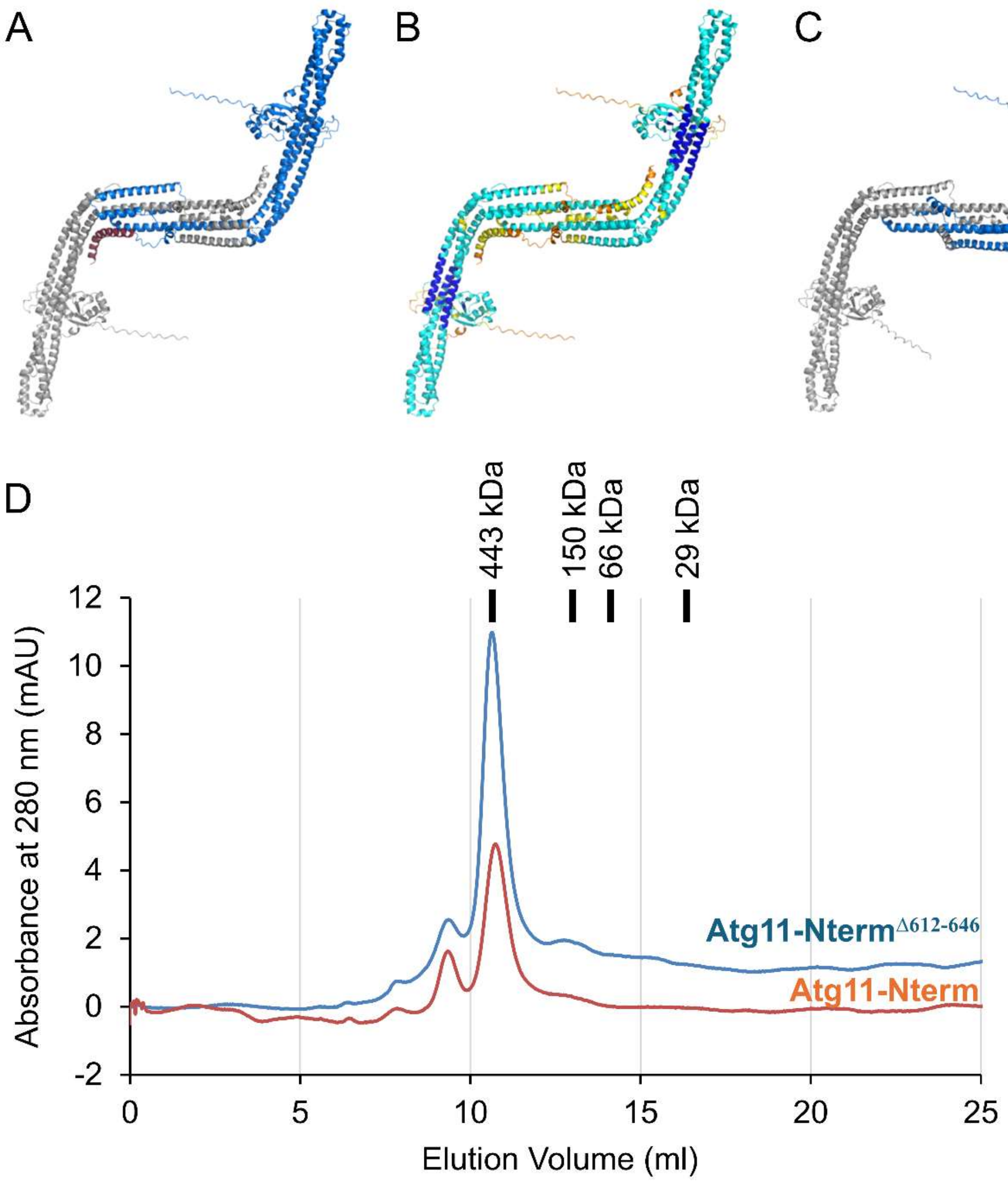


**Supplementary Figure 4. Deletion of a predicted amphipathic helix from Atg11 does not disrupt the structure of the N-terminal region. (A**) AlphaFold 3 model of Atg11-Nterm. One monomer is colored blue and the other monomer is colored grey. The predicted amphipathic helix is colored in dark red for the subunit that is colored blue. (**B**) AlphaFold confidence (pLDDT) scores for Atg11-Nterm. The pLDDT scores are colored dark blue for very high, blue for confident, yellow for low and orange for very low. The majority of Atg11-Nterm has confident or very high pLDDT scores whereas the predicted amphipathic helix is low or very low. (**C**) AlphaFold 3 prediction of Atg11-Nterm$^{\Delta612\text{-}646}$ which lacks the predicted amphipathic helix. (**D**) SEC chromatography of Atg11-Nterm and Atg11-Nterm$^{\Delta612\text{-}646}$ on a Superdex 200 Increase 10/300 GL column in 20 mM Tris pH 8.0, 300 mM NaCl and 0.2 mM TCEP. The molecular weight of four different standards run on the same column in the same buffer are marked above the chromatograms.

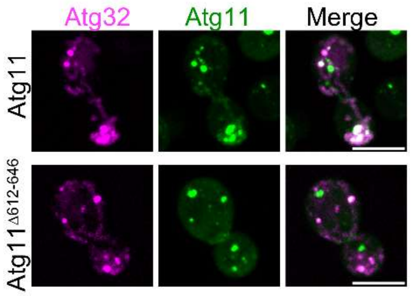


**Supplementary Figure 5.** Atg11Δ612-646 is recruited to Atg32 punctae. Representative images of Δatg11 cells showing 2xGFP-Atg11 or 2xGFP-Atg11Δ612-646 and RFP-Atg32 localization under nitrogen starvation conditions.

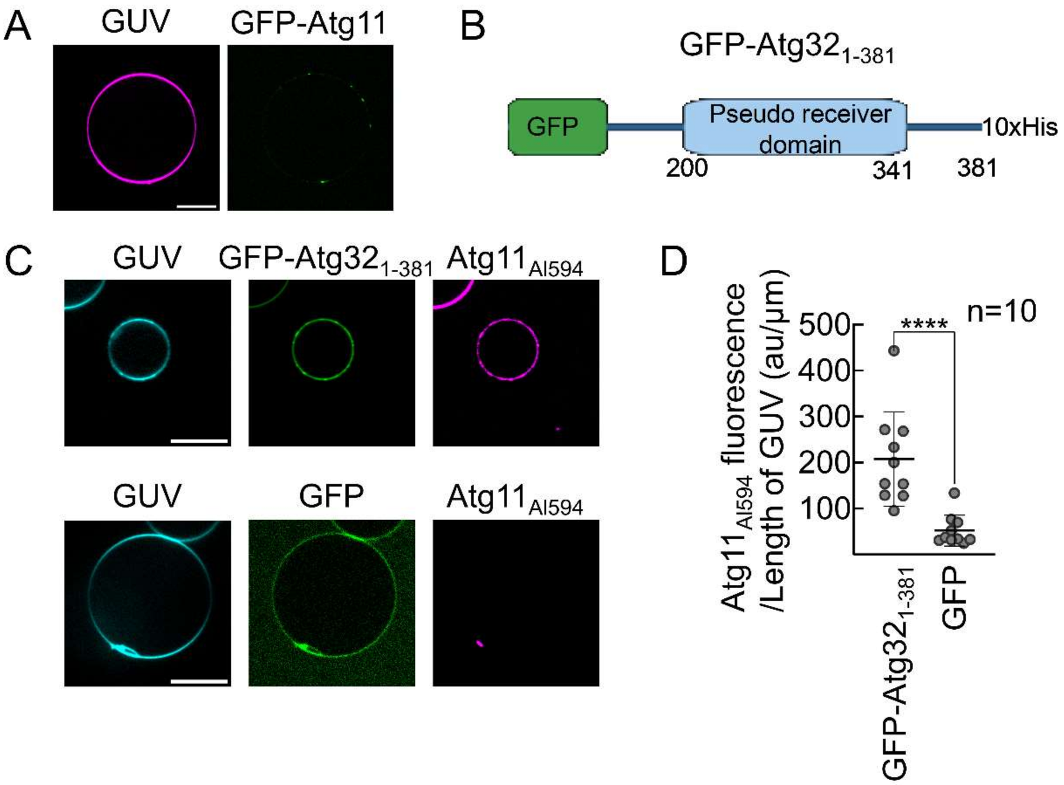


**Supplementary Figure 6. Atg11 binds the N-terminal cytosolic domain of Atg32.** (**A**) Representative fluorescence images of GFP-Atg11 and GUVs mimicking the composition of the outer mitochondrial membrane. (**B**) The domain architecture of $Atg32_{1-381}$ fused to GFP at the N-terminus and containing a 10xHis tag at the C-terminus. GUVs containing 5 mol% of DGS-$Ni^{+2}$NTA were incubated with GFP $Atg32_{1-381}$-10xHis or 6xHis-GFP (control) and mixed with $Atg11_{Al594}$. (**C**) Representative fluorescence images of $Atg11_{Al594}$ binding to GUVs displaying GFP-$Atg32_{1-381}$, but not to GUVs displaying GFP alone. Scale bar=10 µm. (**D**) Quantification of the fluorescence intensity of $Atg11_{Al594}$ along the GUVs normalized to the length of the GUV in presence of GFP-$Atg32_{1-381}$ and GFP. The data are presented as means, with error bars indicating standard deviation. **** $p<0.0001$ calculated using Mann Whitney's test.

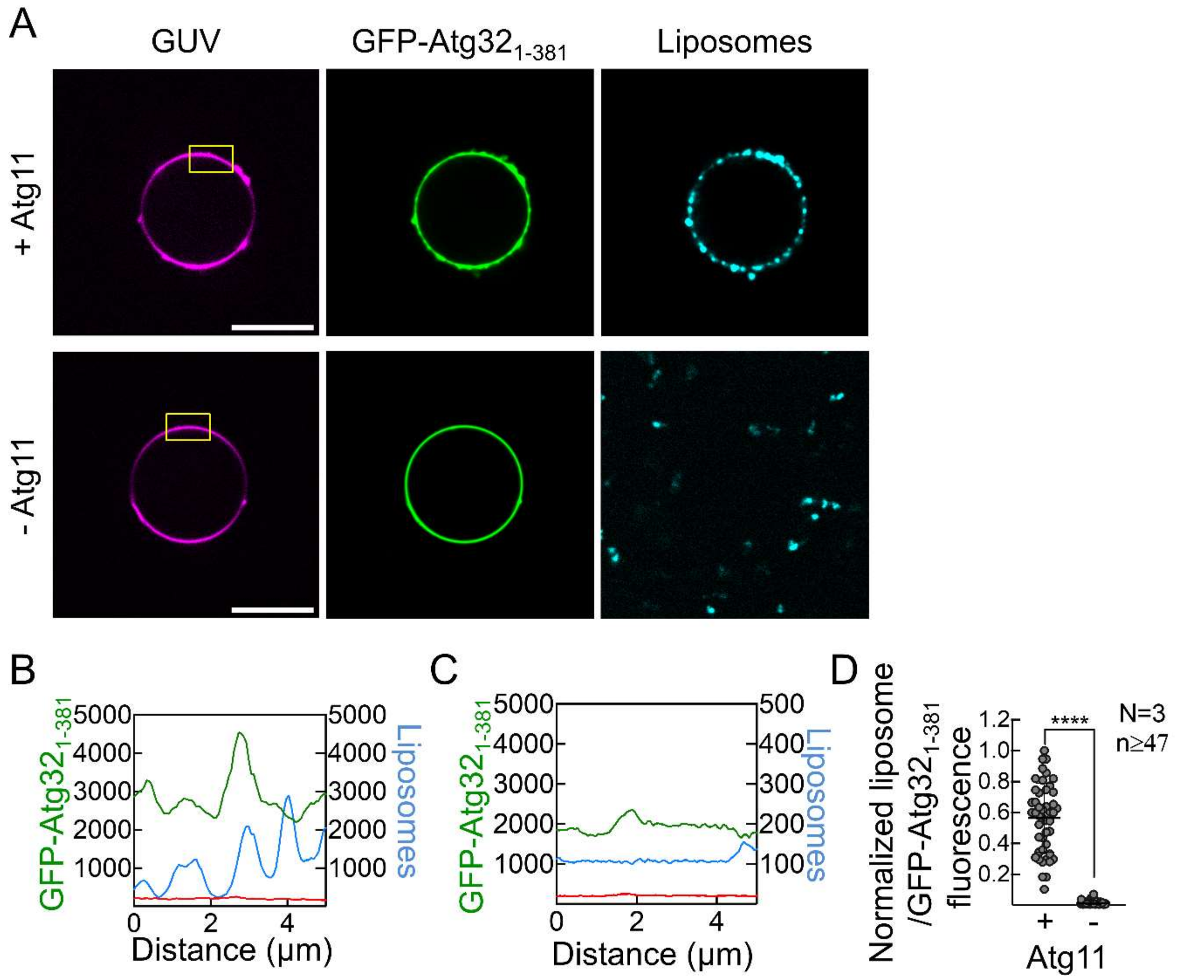


**Supplementary Figure 7. The Atg32-Atg11 complex tethers YPL membranes.** (**A**) Representative fluorescence images of GLT with GUVs containing 5% mol DGS-$Ni^{+2}$NTA displaying His-GFP-Atg32$_{1-381}$ which were sequentially incubated with Atg11 and YPL liposomes (top panel) or YPL liposomes alone (bottom panel). Yellow box marks the region on the GUV used for tracing line profiles in **B** and **C**. Scale bar=10 μm. (**B**) Fluorescence intensity profiles from a line drawn along the GUV in A (top panel) showing liposome and GFP-Atg32$_{1-381}$ fluorescence intensities along the GUV. (**C**) Fluorescence intensity profiles from a line drawn along the GUV in A (bottom panel) showing liposome and GFP-Atg32$_{1-381}$ fluorescence intensities along the GUV. (**D**) Quantification of the ratio of liposomes to GFP-Atg32$_{1-381}$ fluorescence from A. Data from 3 independent experiments (N=3) were pooled and analyzed using non-parametric Mann-Whitney's test. **** indicates $p<0.0001$. n indicates the total number of GUVs analyzed per condition. GUVs were made with DOPC (74.9mol%), DOPS (20mol%), DGS-$Ni^{+2}$NTA (5mol%), and RhPE (0.1mol%). Liposomes were made with YPL (99.9mol%) and DiD (0.1mol%).
